# Arterial effects of anthracycline: structural and inflammatory assessments in non-human primates and lymphoma patients using ^18^F-FDG positron emission tomography

**DOI:** 10.1101/2024.05.30.596741

**Authors:** Stephen Rankin, Caitlin Fountain, Alastair J Gemmell, Daire Quinn, Alasdair Henderson, John McClure, Sandy Small, Balaji Venugopal, Pamela McKay, Piotr J Slomka, David Colville, Mark C Petrie, Giselle C. Meléndez, Ninian N Lang

**Affiliations:** BHF Glasgow Cardiovascular Research Centre, School of Cardiovascular and Metabolic Health, University of Glasgow, Glasgow UK; Departments of Internal Medicine, Section on Cardiology and Pathology, Section on Comparative Medicine. Wake Forest University School of Medicine, Winston-Salem, USA; Department of Clinical Physics & Bioengineering, NHS Greater Glasgow & Clyde, Glasgow; The Beatson West of Scotland Cancer Centre, Glasgow, UK; Cedars-Sinai, Division of Artificial Intelligence in Medicine, Department of Medicine, Los Angeles, USA; School of Medicine, Dentistry and Nursing, University of Glasgow

**Keywords:** anthracycline, vascular toxicity, fibrosis, PET, lymphoma, doxorubicin

## Abstract

**Background:** Anthracyclines, such as doxorubicin, are important anti-cancer therapies but are associated with arterial injury. Histopathological insights have been limited to small animal models and the role of inflammation in the arterial toxic effects of anthracycline is unclear in humans. Our aims were: 1) To evaluate aortic media fibrosis and injury in non-human primates treated with anthracyclines; 2) To assess the effect of anthracycline on aortic inflammation in patients treated for lymphoma.

**Methods:** 1) African Green monkeys (AGM) received doxorubicin (30–60 mg/m^2^/biweekly IV, cumulative dose: 240 mg/m^2^). Blinded histopathologic analyses of collagen deposition and cell vacuolization in the ascending aorta were performed 15 weeks after the last doxorubicin dose and compared to 5 age- and gender-matched healthy, untreated AGMs. 2) Analysis of the thoracic aorta of patients with diffuse large B-cell lymphoma (DLBCL), at baseline and after doxorubicin exposure, was performed using ^18^F-fluorodeoxyglucose (^18^F-FDG) positron emission tomography/computed tomography (PET/CT) in this observational study. The primary outcome was change in maximal tissue-to-background ratio (TBRmax) of the thoracic aorta from baseline to their end-of-treatment clinical PET/CT.

**Results:** In AGMs, doxorubicin exposure was associated with greater aortic fibrosis (collagen deposition: doxorubicin cohort 6.23±0.88% vs. controls 4.67±0.54%; p=0.01) and increased intracellular vacuolization (doxorubicin 66.3 ± 10.1 vs controls 11.5 ± 4.2 vacuoles/field, p<0.0001) than untreated controls.

In 101 patients with DLBCL, there was no change in aortic TBRmax after anthracycline exposure (pre-doxorubicin TBRmax 1.46±0.16 vs post-doxorubicin TBRmax 1.44±0.14, p=0.14). The absence of change in TBRmax was consistent across all univariate analyses.

**Conclusions:** In a large animal model, anthracycline exposure was associated with aortic fibrosis. In patients with lymphoma, anthracycline exposure was not associated with aortic inflammation.Further research is required to elucidate the mechanisms of anthracycline-related vascular harm.

## INTRODUCTION

Anthracyclines, such as doxorubicin, are effective anti-cancer drugs used as the backbone of treatment of numerous cancer types, including breast cancer, sarcoma, and lymphoma. However, these chemotherapeutic agents are associated with cardiovascular toxicities including arterial and myocardial injury.^1^ It is hypothesized that coronary endothelial injury may contribute to the development of left ventricular dysfunction and heart failure while peripheral arterial injury may further amplify these risks via arterial stiffening and consequent disruption of ventriculo-arterial coupling ^2^. While aortic stiffening has been observed in humans after exposure to anthracycline, assessment of the underlying pathophysiologic mechanisms has been limited to small animal and cell line studies ^3,4^.

The role of inflammation as a mediator of vascular dysfunction is well recognized^5–7^and is a notable target for the prevention or treatment of a range of arterial diseases and injurious processes.^8–10^ Furthermore, in small animal models exposed to anthracyclines, inflammation-associated aortic stiffening^11^ and peri-vascular inflammation have been demonstrated.^12^ While these observations raise the hypothesis that anthracyclines induce inflammation-mediated arterial injury in human patients, this has not yet been established. ^18^Fluoride-fluorodeoxyglucose positron emission tomography/computed tomography (^18^F-FDG PET/CT) is a molecular imaging technique that is highly sensitive to metabolically active processes that use glucose as a fuel. It is in routine clinical use for the staging of a range of cancers and the assessment of treatment responses. Notably, it is also the gold standard method for the identification and quantification of inflammatory activity, including in large arteries^13–16^.

In this study, we examined the histopathological and inflammatory effects of exposure to doxorubicin in a non-human primate model and in patients before and after chemotherapy for the treatment of lymphoma. We evaluated these effects in order to inform detection, prevention and treatment strategies for patients at risk of anthracycline-associated arterial toxicity.

## METHODS

### Non-human Primate Study

This non-human primate study (NHP) was performed in accordance with the *Guide for Care and Use of Laboratory Animals* and approved by the *Wake Forest School of Medicine Institutional Animal Care and Use Committee*. The institution is accredited by the *Association for the Assessment and Accreditation of Laboratory Animal Care International* and operates in compliance with the Animal Welfare Act.

Five female pre-menopausal African Green monkeys (AGM) (*Chlorocebus aethiops sabeus*) aged 13 ± 1.3 years were used in this study. The study subjects were sourced from a multigenerational pedigreed colony of African Green monkeys (n=311, 4-27 years, lifespan ≈ 26 years), which descended from 57 founder monkeys at the Wake Forest Vervet Research Colony (P40-OD010965). Animals were fed a commercial laboratory primate chow (Laboratory Diet 5038; LabDiet, St. Louis, MO) with daily supplemental fresh fruits and vegetables and tap water ad libitum. Monkeys were housed in a climate-controlled room. As previously described ^17,18^, animals underwent doxorubicin treatment which consisted of two initial doses of 30 mg/m^2^ and three doses of 60 mg/m^2^ given via vascular access port (VAP) every 17 ± 3.5 days (total cumulative dose: 240 mg/m^2^). At the experimental endpoint (15 weeks after the last dose of doxorubicin), euthanasia was induced in accordance with *American Veterinary Medical Association (AVMA)* guidelines. While under anesthesia, a catheter was placed in a peripheral vein, and euthanasia solution was administered at a dose of sodium pentobarbital ∼100 mg/kg IV. Euthanasia was achieved by exsanguination and subsequent removal of the heart. Ascending aorta cross-sections were acquired from ∼1 cm above the sinotubular junction and fixed in 4% paraformaldehyde for subsequent histopathologic analysis. For non-treated control samples, archival tissues from 5 age- and sex-matched healthy animals were used for histopathological comparisons.

#### Histopathology

Histological images of the aortas were captured using a digital slide scanner (Hamamatsu HT NanoZoomer). Five μm consecutive sections from each block were stained with hematoxylin and eosin (H&E) and Masson’s trichrome staining. Tunica media collagen volume fraction (CVF) and vacuoles were quantified from photomicrographs of 10 random fields from each aortic section obtained using a 20X objective^19^. Collagen volume fraction was determined from these images using Image J (NIH, Bethesda, MA) and expressed as a percentage of area. The number of vacuoles from each microphotograph was annotated and expressed as the mean of the 10 microscopy fields. The aortic medial area was measured in low magnification H&E-stained sections by tracing the circumference of the external and internal elastic lamina; the area was calculated by subtracting the area of the internal from the external circumference. Medial thickness was determined by measuring the distance from the internal to external elastic lamina from four random points across the aorta. All analysis was performed by researchers blinded to the treatment group.

### Clinical Study in Patients with Lymphoma

Ethical approval was granted by the West of Scotland Research Ethics Committee (22/WS/0180). This was a retrospective observational study with no change in patient management. Prior to each clinical PET scan, patients were asked to consent for images to be used in future research as part of the routine clinical pre-PET questionnaire. Those who consented were included. All aspects of this study were performed in accordance with the Declaration of Helsinki.

We performed a retrospective review of ^18^F-FDG-PET/CT scans of patients with diffuse large B-cell lymphoma (DLBCL) referred to the Regional PET service at Glasgow Beatson West of Scotland Cancer Centre between 2019 to 2023. This is a cancer network hub center which receives referrals from 16 referring hospital sites. We included patients aged 18 years and older treated using anthracycline regimens receiving a cumulative dose of >150mg/m^2^ of doxorubicin. We excluded patients whose arterial FDG uptake could not be quantified due to significant interference from adjacent lymphoma. Other exclusion criteria were those treated at a hospital that was not in the NHS Greater Glasgow & Clyde catchment area, incomplete medical records, concurrent thoracic radiotherapy, technically inadequate scans (such as non-diagnostic tracer uptake), blood glucose >11mmol/L before either scan and suspected or confirmed vasculitis on the baseline scan. Patients were also excluded if their baseline PET/CT was more than 3 months before starting chemotherapy.

Detailed baseline demographic data, including past medical history and cancer history, were collected from electronic case note reviews. Baseline cardiovascular risk stratification was performed using the European Society of Cardiology (ESC) baseline cardio-oncology CV risk stratification assessment tool^20^. Response to treatment was collected by the Deauville score reported on the clinical scan.

#### ^18^F-FDG PET/CT imaging

^18^F-FDG-PET/CT scanning was performed in accordance with departmental standard procedures based on *European Association of Nuclear Medicine (EANM)* guidelines ^21^ on PET/CT scanners (Discovery-690 or 710, General Electric System, Milwaukee, WI, USA or Biograph Vision 600, Siemens, Erlangen, Germany). Patients were fasted for a minimum of 6 hours prior and blood glucose levels were checked during patient preparation. Scanning was performed 1 hour after intravenous administration of 4 MBq/kg ^18^F-FDG. CT images were acquired at 120kV, with automatic mA modulation applied (Noise Index=30 or reference mAs=50, dependent on the scanner used) and covered from the base of the skull to mid-thigh, reconstructed at 1.5-2.5mm increments. PET images encompassed the same transverse field of view as the CT. PET acquisition times were 3-4 min per bed position or 1-1.5mm/second, depending on the scanner used. PET attenuation correction was based on CT, and images were corrected for the scatter, time-of-flight and point spread function, and iteratively reconstructed using local clinical reconstruction parameters.

#### PET analysis

Analysis of aortic ^18^F-FDG uptake was performed using FusionQuant v1.21 software (Cedars-Sinai Medical Centre, Los Angeles). For all methods, co-registration between PET signal and CT images was ensured in 3 orthogonal planes. Care was taken to ensure activity from adjacent tissue, such as esophagus or adjacent lymphoma, was not included in the aortic analysis. Background activity in the blood pool was determined as the mean standardized uptake value (SUVmean) of 10 sequential cylindrical 3-Dimensional (3D) volumes of interest (VOI) within the superior vena cava (SVC), starting at the confluence of the innominate vein. Aortic ^18^F-FDG activity was measured using 4mm thick 3D polygonal VOIs starting at the ascending aorta to the end of the thoracic descending aorta. Aortic analysis started at the inferior aspect of the right pulmonary artery and stopped when the descending thoracic aorta passed through the diaphragm. FDG uptake of the arterial vessel compared to background uptake was used, giving a target-to-background ratio (TBR), in accordance with EANM guidelines^13^.

TBR was calculated for each VOI by dividing the SUVmax and SUVmean value by the blood pool activity to give TBRmax and TBRmean, respectively. TBR values were then averaged for the thoracic aorta and each aortic segment. Analysis was performed in concordance with EANM recommendation for aortic activity including TBRmax, TBRmean, activity within ‘active segments’ (defined as a TBRmax ≥1.6) and most diseased segment (MDS, the three consecutive VOIs around the VOI with the highest activity to represent the most intense lesion), described previously^13^. 10% of scans were randomly selected and re-analyzed by two trained observers (SR, DC) for inter-observer and intra-observer agreement. To minimize recall bias, intra-observer repeatability was assessed by the same trained researcher (SR) using repeated assessments performed 3 months apart in random order.

#### Calcium scoring

Aortic calcification assessment was performed on the CT of the baseline PET/CT scan on a dedicated workstation (Vitrea Advanced, Vital Imaging, Toshiba Systems, Minnesota, USA). A density threshold of 130 Hounsfield units, 3-pixel threshold on 3mm slice thickness was used and a cumulative calcium score of the whole aorta and each thoracic aortic segment was calculated as previously described ^22^.

#### Statistical analysis

Statistical analysis was performed using STATA software (Version 17). Continuous data with normal distribution are presented as mean ± standard deviation (SD), and skewed data are presented as median and interquartile range (IQR). Between groups comparisons were made using paired t tests and ANOVA or non-parametric equivalents as appropriate. Normal distribution was assessed by the Shapiro-Wilk test. Intra- and interobserver variability was assessed by the intra-class correlation (ICC) coefficient. Based on prior studies using change in mean TBRmax as a primary outcome, a sample size of 101 patients was needed to detect a 10-15% difference between groups with a power of 85% and significance of 5%, assuming a baseline TBRmax of 1.57 ± 0.42 and a SD of change in TBRmax of 0.4^23–25^. A *p* value <0.05 was taken to represent statistical significance.

## RESULTS

### Non-Human Primate Study

Both groups were of similar age (control 12.5 ± 0.7 vs doxorubicin 13.1 ± 0.6y, p=0.3), weight (control: 5.4 ± 0.5 vs doxorubicin: 4.9 ± 0.3kg, p=0.2), and body surface area (BSA, control: 0.28 ± 0.01 vs doxorubicin: 0.27 ± 0.01, p=0.2). Morphometric data of the animals have been previously described ^17,18^.

Histopathological analysis of the structural changes within hematoxylin-eosin stained aortas of AGMs exposed to doxorubicin there was more intracellular vacuolization in comparison to control animals (control 11.5 ± 4.2 vs doxorubicin 66.3 ± 10.1 vacuoles/field, p<0.0001, Figure 1). Doxorubicin treatment was associated with an increase in collagen deposition in the aortic media consistent with fibrosis (controls: 4.67 ± 0.54% vs. doxorubicin: 6.23 ± 0.88%, p=0.01; Figure 2). There was no difference in the medial area (controls: 10.6 ± 2.03mm^2^ vs dox: 9.32 ± 2.27mm^2^, p=0.3) nor medial area/BSA (control: 2200 ± 404.3 mm^2^/m^2^ vs doxorubicin: 2127 ± 462.2 mm^2^/m^2^, p=0.7) between groups and medial thickness was also not different between control animals and those exposed to doxorubicin (control: 573.2 ± 71.61μm vs doxorubicin: 519.4 ± 96.15 μm, p=0.34).

**Figure 1.**
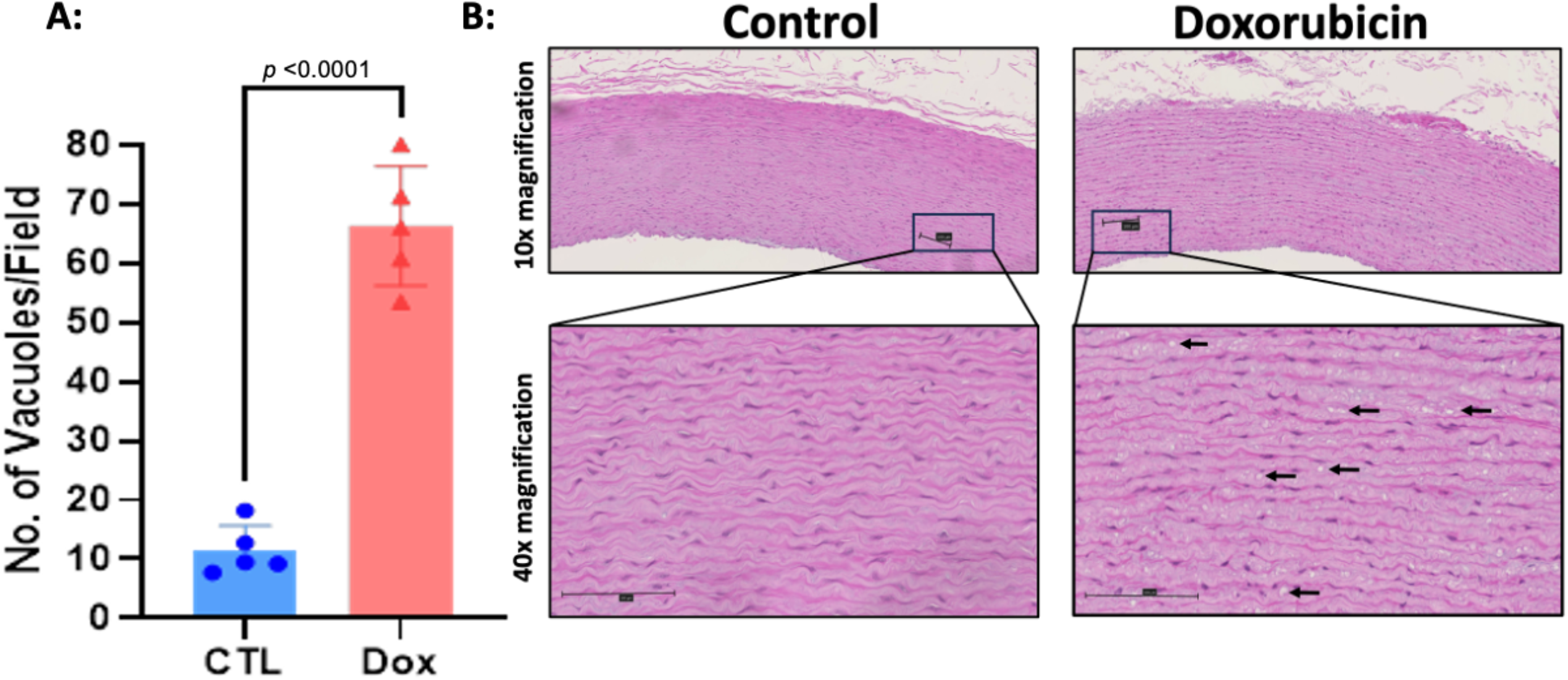
Intracellular vacuolization within the aortic media of monkeys exposed to doxorubicin compared with controls. *Histopathological assessment of hematoxylin-eosin stained aortas. A: Graphical representation of the number of vacuoles present in the arterial wall in the control arm (CTL) and the doxorubicin treated arm (Dox), p<0.0001. (B). Representative microphotographs of hematoxylin-eosin stained aortas at 10X and 40X magnification. Black arrows indicate cardiomyocytes with intracellular vacuolization. All values are mean ± SEM*.

**Figure 2.**
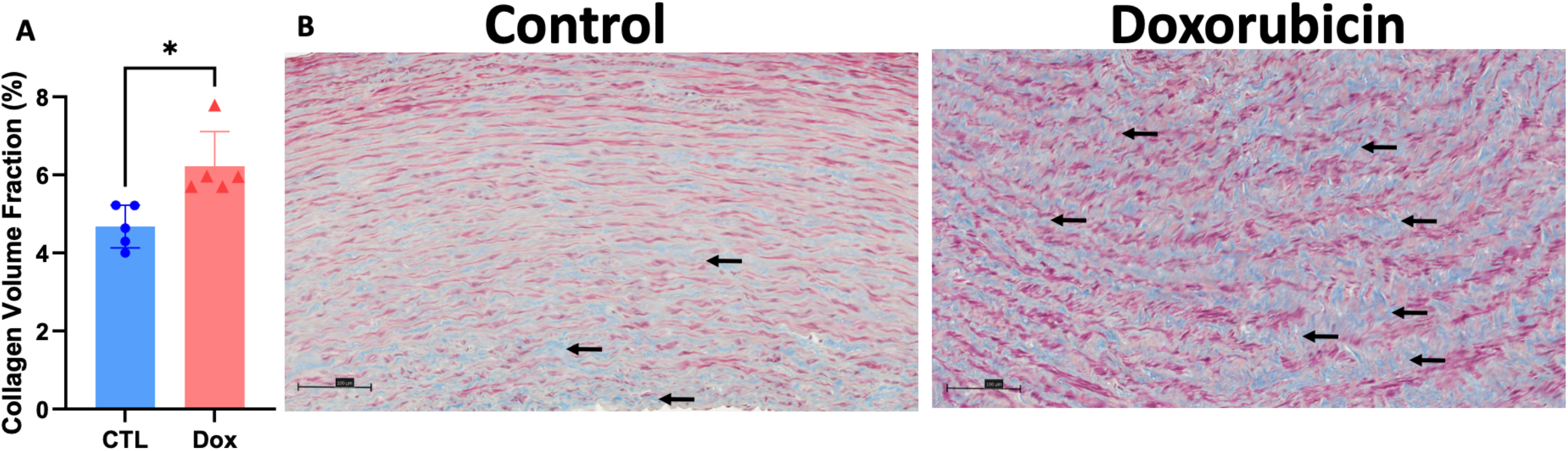
Collagen fiber deposition in the aortic media of monkeys exposed to doxorubicin compared with controls. *Assessment of ascending aorta media deposition of interstitial collagen. Graphical representation of collagen volume fraction (A) in aortas of untreated controls (blue bar, n=5) and Dox-treated AGMs (red bar, n=5). Representative microphotographs (20X) of aortas of untreated controls (B) and Dox-treated AGMs(C). Collagen is stained blue (highlighted by black arrows), and the cytoplasm of smooth muscle cells is stained red and pink in the microphotographs. All values are mean ± SEM. * p=0.01*

### Clinical Study in Patients with Lymphoma

A total of 101 patients were included in the analysis (Supplementary Figure 1) of whom the mean age was 64 ± 12 years and 47 were female. The majority of patients had advanced lymphoma (71% with stage III/IV disease). The median cumulative doxorubicin dose was 300mg/m^2^ (IQR 225-300mg/m^2^) and 82% of patients received 6 cycles of doxorubicin. There was a high prevalence of cardiovascular risk factors (hypertension, dyslipidemia, ischemic heart disease, smoking history, diabetes, body mass index ≥35kg/m^2^) with 69% of patients having at least one. On the basis of ESC Cardio-Oncology cardiovascular risk stratification criteria, 55% of patients would be considered to have at least ‘medium’ cardiovascular risk (36% ‘medium’, 18% ‘high’ and 1% ‘very high’ risk, Table 1). The median time from baseline PET/CT to starting doxorubicin was 7 days (IQR 4-14 days). The mean time between starting doxorubicin and post-doxorubicin PET/CT was 4.8±0.9 months and the median time from finishing chemotherapy to follow up PET/CT was 1.2 months (IQR 0.9-1.6 months).

**Table 1:** Baseline demographics of patients with diffuse large B-cell lymphoma (DLBCL), treated with anthracycline-based chemotherapy.

| Demographics |  | Total (n=101) |
| --- | --- | --- |
| Mean age (years ± SD) | | 64.3±12 |
| Sex (n/%) | Male | 54 (54) |
|  | Female | 47 (47) |
| Mean BMI (kg/m<sup>2</sup> ± SD) | | 27.8±5.8 |
| Hematological characteristics |  |  |
| Cancer stage (n/%) | 1 | 14 (14) |
|  | 2 | 16 (16) |
|  | 3 | 24 (24) |
|  | 4 | 47 (47) |
| Previous chemotherapy (n/%) | No | 86 (86) |
|  | Non-anthracycline regimen | 10 (10) |
|  | Anthracycline regimen | 4 (4) |
| Anthracycline regimen (n/%) | RCHOP | 96 (95) |
|  | R-CODOX-M-IVAC | 5 (5) |
| Cycles of anthracycline (n/%) | $\leq 3$ | 12 (12) |
|  | 4-5 | 5 (5) |
| | $\geq 6$ | 84 (84) |
| Median doxorubicin cumulative dose (mg/m<sup>2</sup>) |  | 300.0 (IQR 225.0-300.0) |
| Baseline cardiovascular characteristics |  |  |
| ESC Baseline CV risk stratification (n/%) | Low risk | 46 (46) |
|  | Medium risk | 36 (36) |
|  | High risk | 18 (18) |
|  | Very high risk | 1 (1) |
| Aortic calcification at baseline (n/%) | Yes | 78 (78) |
|  | No | 23 (23) |
| Median aortic calcium score |  | 213.0 (IQR 11.0-938.0) |
| Previous myocardial infarction (n/%) |  | 5 (5) |
| Previous coronary revascularization* (n/%) |  | 3 (3) |
| Angina (n/%) |  | 2 (2) |
| Previous stroke (n/%) |  | 7 (7) |
| Previous heart failure (n/%) |  | 1 (1) |
| History of atrial fibrillation (n/%) |  | 3 (3) |
| Hypertension (n/%) |  | 35 (35) |
| Diabetes (n/%) |  | 14 (14) |
| Smoking history (n/%) | ex-smoker | 27 (27) |
|  | current | 18 (18) |
| Antiplatelet therapy (n/%) |  | 12 (12) |
| ACEI or ARB (n/%) |  | 28 (28) |
| Diuretic therapy (n/%) |  | 5 (5) |
| Beta-blocker (n/%) |  | 18 (18) |
| Rate limiting CCB (n/%) |  | 3 (3) |
| Non-rate limiting CCB (n/%) |  | 12 (12) |
| Statin (n/%) |  | 26 (26) |
| ACEI – Angiotensin-converting enzyme inhibitor, ARB – angiotensin receptor blocker, BMI- Body Mass Index, CCB – calcium channel blocker, CV- cardiovascular, RCHOP- rituximab, cyclophosphamide, doxorubicin, vincristine, prednisolone. R-CODOX-M-IVAC- rituximab, cyclophosphamide, cytarabine, vincristine, doxorubicin, methotrexate, ifosfamide, etoposide, cytarabine *prior percutaneous coronary intervention or coronary artery bypass grafting. |  |  |

#### PET analysis

The mean TBRmax of the thoracic aorta at baseline and follow up was 1.46 ± 0.16 vs 1.43 ± 0.14, respectively, p=0.14, Figure 3. In comparison to baseline, there was no observed difference in aortic inflammation, measured by mean TBRmax, TBRmean, TBRmax within ‘active segments’ or ‘most diseased segment (MDS)’ after doxorubicin exposure, Table 2. Comparison of ^18^F-FDG uptake by each aortic segment was similar to that of the whole aorta before and after doxorubicin. There was a very small decrease in mean TBRmax after doxorubicin observed within the aortic arch that was of borderline statistical significance (mean TBRmax pre-doxorubicin: 1.52 vs post-doxorubicin: 1.49; - 0.03 difference, 95% CI -0.65-0, p*=*0.05), Table 2.

**Table 2:** Aortic TBR and MDS measured by PET, before and after anthracycline-based chemotherapy.

| Aortic Segment | PET parameter | Mean TBRmax |  | Difference | 95% CI | *p* |
| --- | --- | --- | --- | --- | --- | --- |
|  |  | Pre-dox | Post-dox |  |  |  |
| Whole aorta | TBRmax | 1.46 ±0.16 | 1.44 ±0.14 | -0.02 ±0.15 | -0.05 to 0.01 | 0.14 |
|  | TBRmean | 1.06 ±0.10 | 1.05 ±0.09 | -0.01 ±0.1 | -0.03 to 0.01 | 0.29 |
|  | TBR max of active segments | 1.71 ±0.08 | 1.7 ±0.06 | -0.01 ±0.08 | -0.03 to 0.07 | 0.18 |
|  | MDS | 1.65 ±0.22 | 1.64 ±0.20 | -0.01±0.23 | -0.06 to 0.03 | 0.62 |
| Ascending | TBRmax | 1.51 ±0.18 | 1.48 ±0.15 | -0.02 ±0.15 | -0.05 to 0.01 | 0.19 |
|  | TBRmean | 1.07 ±0.11 | 1.06 ±0.09 | -0.01 ±0.1 | -0.03 to 0.07 | 0.19 |
|  | TBR max of active segments | 1.74 ±0.11 | 1.70 ±0.07 | -0.03 ±0.11 | -0.07 to 0.04 | 0.08 |
|  | MDS | 1.57 ±0.19 | 1.55 ±0.17 | -0.02 ±0.18 | -0.06 to 0.01 | 0.20 |
| Arch | TBRmax | 1.52 ±0.16 | 1.49 ±0.14 | -0.03 ±0.16 | -0.65 to 0.001 | 0.05 |
|  | TBRmean | 1.07 ±0.08 | 1.06 ±0.07 | -0.01 ±0.08 | -0.02 to 0.09 | 0.36 |
|  | TBR max of active segments | 1.75 ±0.10 | 1.71 ±0.12 | -0.03 ±0.15 | -0.08 to 0.02 | 0.18 |
|  | MDS | 1.61 ±0.20 | 1.57 ±0.18 | -0.04 ±0.22 | -0.08 to 0.00 | 0.06 |
| Descending | TBRmax | 1.43 ±0.16 | 1.41 ±0.15 | -0.02 ±0.16 | -0.05 to 0.01 | 0.21 |
|  | TBRmean | 1.05 ±0.12 | 1.04 ±0.11 | -0.01 ±0.11 | -0.03 to 0.01 | 0.33 |
|  | TBR max of active segments | 1.70 ±0.08 | 1.70 ±0.07 | -0.002 ±0.1 | -0.03 to 0.03 | 0.88 |
|  | MDS | 1.60 ±0.20 | 1.59 ±0.22 | -0.01 ±0.24 | -0.05 to 0.04 | 0.82 |

**Figure 3.**
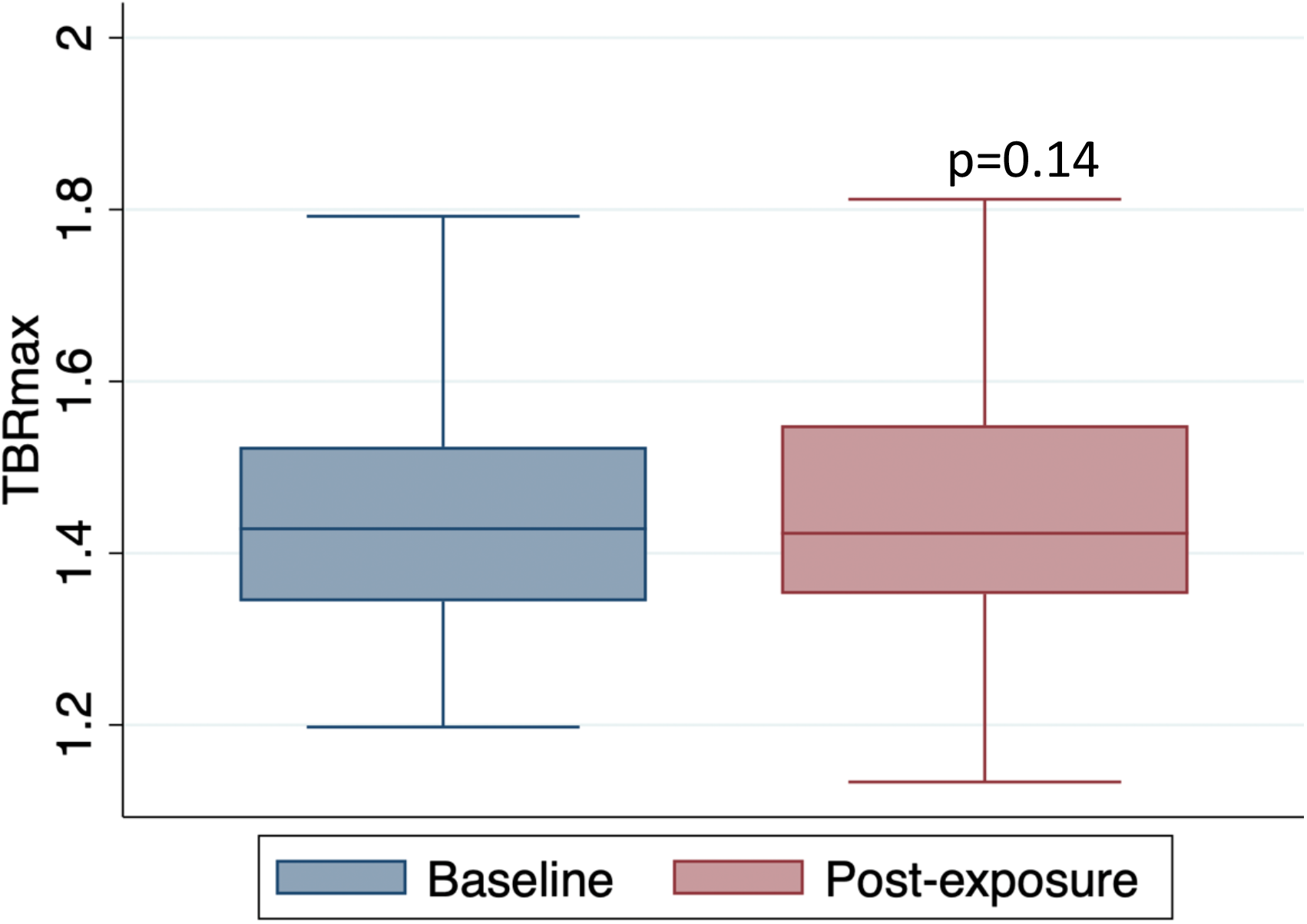
FDG uptake of the thoracic aorta in patients with lymphoma before and after treatment with anthracycline-based chemotherapy. *Boxplot of the mean TBRmax of the whole aorta pre and post anthracycline exposure, p=0.14*.

**Figure 4.**
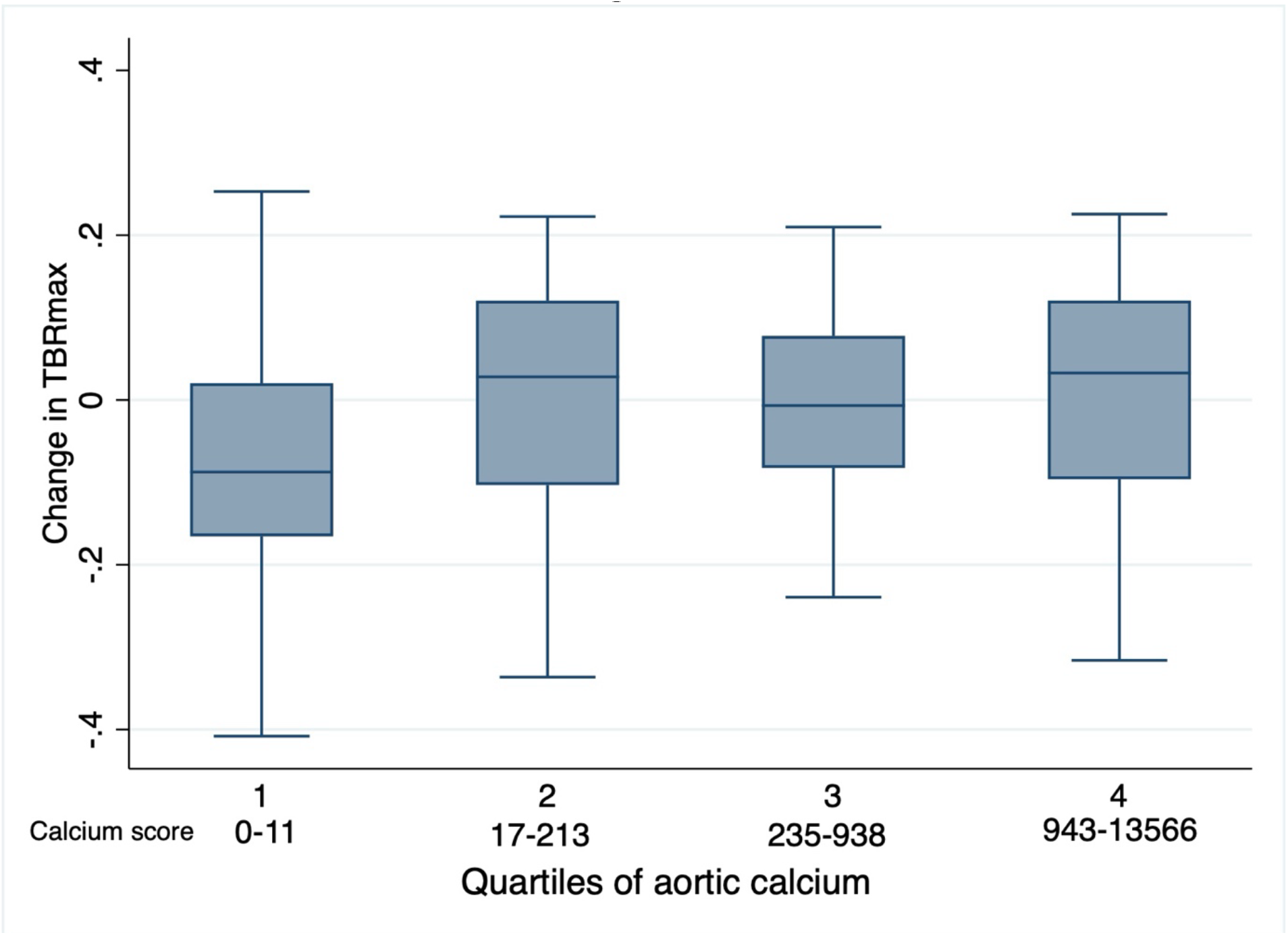
Change in mean TBRmax y baseline aortic calcium score in patients with lymphoma before and after treatment with anthracycline-based chemotherapy. *The change in aortic TBRmax from before and after anthracycline compared with quartiles of aortic calcification, p=0.42*.

Intra-observer and inter-observer assessments were highly correlated and demonstrated excellent reproducibility (Supplementary Table 1).

#### Baseline aortic Calcification & ^18^F-FDG uptake

Aortic calcification was present on the baseline PET/CT scan in 78 of the 101 patients (calcium score >0, Table 1). Aortic calcium scores ranged from 0 to 13566, with a median score of 213 (IQR 11-938). TBRmax remained unchanged from baseline, irrespective of the baseline aortic calcium score: there was no difference in TBRmax when comparing patients grouped by quartile of baseline aortic calcification, including comparison of the lowest quartile (aortic calcium score range 0-11) with the highest quartile (calcium score range 943-13566), p=0.42.

#### PET analysis by clinical factors

Univariate analysis of baseline demographics and cardiovascular risk factors (hypertension, dyslipidemia, ischemic heart disease, smoking history, diabetes, body mass index ≥35kg/m^2^) did not identify any association between these variables with change in aortic FDG uptake, when assessed by TBRmax, TBRmean, MDS and ‘active segments,’ after doxorubcin exposure. Baseline CV risk, assessed using the ESC risk stratification tool, was also not associated with change in aortic FDG uptake (Supplementary Table 2). There was no association between the change in aortic activity and cumulative doxorubicin dose, stage of disease at diagnosis, or response to treatment on follow up imaging (Supplementary Table 3).

## DISCUSSION

We investigated the arterial effects of anthracyclines in a non-human primate model and in patients with lymphoma. Doxorubicin, a very commonly used anthracycline drug, was associated with substantial aortic vacuolization and fibrosis in monkeys. In contrast, we did not observe more inflammation in the thoracic aorta of patients exposed to doxorubicin in patients treated for lymphoma. ^18^F-FDG PET/CT, the gold standard imaging modality for arterial inflammation, was used and did not identify inflammation as a pathogenetic mechanism for the anthracycline related fibrosis.

Our non-human primate study provides novel insights into arterial morphologic changes and extracellular matrix remodeling induced by anthracyclines. By employing this unique large animal model that shares similar genetic traits with humans, our results provide valuable and potentially translatable data. Our animal study incorporated a doxorubicin dosing scheme similar to that received by the patients in our clinical study. Anthracycline exposure was associated with substantial histopathological vascular changes including greater deposition of vascular collagen and intracellular vacuolization of the arterial media. We have previously reported that these animals developed cardiac fibrosis and an absolute reduction of left ventricular (LV) ejection fraction of 25% ^18^. However, to the best of our knowledge, the effects of anthracyclines on arterial remodeling have not previously been explored in a large animal model. Small animal models consistently demonstrate that inflammation is implicated in the pathophysiology of hypertension- and age-related vascular dysfunction^26,27^ and there are overlaps between the structural and functional consequences of these pathologies and anthracycline-associated arterial toxicity, including extracellular matrix remodeling, degradation of elastin, and formation of advanced glycated end products ^3,4,26,27^. In animal models of age-related vascular dysfunction, pro-inflammatory pathways are active, including tumour necrosis factor-alpha (TNF-α), interleukins-1β and -6, with macrophage and T cell infiltration observed in the adventitia and surrounding adipose tissue ^28,29^. Similar circulating biomarkers, and in mouse aortic lysates, have been observed in small mouse models of anthracycline vascular toxicity^11^. It is also plausible that arterial inflammation also occurs in the coronary macro- and microvasculature to contribute to myocardial toxic effects of anthracycline. Notably, prior work has demonstrated a link between cardiomyocyte vacuolization and tissue edema ^30^ while we and others have demonstrated that cardiac fibrosis is preceded by inflammation and edema ^18,30,31^. Intriguingly, the recent STOP-CA trial demonstrated that atorvastatin prevents anthracycline-associated cardiotoxicity in patients with lymphoma^32^. The mechanism underlying this effect has not been defined but it is possible that arterial protective effects, including anti-inflammatory, may be at least partially responsible.

In humans, prior treatment with anthracycline is associated with elevated arterial stiffness and this may be a consequence of arterial fibrosis^33^. Elevated arterial stiffness exacerbates ventriculo-arterial uncoupling which, in turn, contributes to LV pressure overload, adverse remodeling and LV dysfunction. Therefore, understanding the mechanisms via which anthracycline evokes arterial fibrosis and stiffening is of fundamental importance so that patients can receive optimal cancer treatment while preventative strategies are developed to minimize adverse cardiovascular effects.

In light of our pre-clinical data, we hypothesized that inflammation may be central to the development of anthracycline-associated arterial stiffening in humans. We assessed aortic inflammation in a cohort of 101 patients with lymphoma treated with anthracycline. Capitalizing upon clinical datasets, we had detailed cardiovascular and oncologic information available. We excluded patients with low-anthracycline dose exposures and the median cumulative doxorubicin dose of 300 mg/m^2^ received by these patients was similar to that used in the non-human primate study. The patients included were representative of the ‘real world’ population of patients with a mean age of 64 years and an almost equal representation of men and women. Cardiovascular disease and risk factors were prevalent at baseline, with over half of patients considered to be at least moderate risk when assessed using the ESC baseline CV risk stratification assessment tool. Despite these methodological strengths, we observed no change in aortic inflammation when assessed just over a month after the completion of treatment with anthracycline-based chemotherapy. Our primary measure of ^18^F-FDG-PET/CT inflammatory activity was mean TBRmax but the lack of change in inflammatory activity also held true when assessed via other analysis methods including TBRmean, TBRmax within ‘active segments’ and within ‘most diseased segments (MDS)’. There was also no association between inflammatory activity and potential hematologic and cardiovascular risk factors, including aortic calcification in 78% of patients, which we used as an objective and quantifiable marker of pre-existing cardiovascular disease.

We assessed the inflammatory effects of anthracycline around one month after completion of anthracycline. Aortic stiffening has been demonstrated to occur within 4 months of anthracycline exposure and we wished to examine this ‘high risk’ period ^34^. A previous smaller retrospective study examined arterial ^18^F-FDG PET/CT in 52 patients following anthracycline treatment for Hodgkin lymphoma at a mean of 65 weeks and found no difference in large artery TBR ^24^. The longer time between completion of chemotherapy and PET scanning meant that evidence of ongoing active inflammation would have been considerably less likely. Furthermore, in contrast to our study, the mean age of the patients was only 35 years, with very few cardiovascular risk factors and, of particular relevance to immune-related analyses, 35% were HIV positive. Overall, our findings imply that anthracycline-associated arterial fibrosis and stiffening occurs independently of inflammation.

### Limitations

While the animals in the non-human primate study were age- and gender-matched between groups, these animals would be considered to be otherwise healthy and without pre-existing cardiovascular disease or risk factors. Furthermore, although the animals were exposed to clinically relevant doses of anthracycline (as well as using the same agent as received by the patients), they were free of cancer and a potential interaction between anthracycline exposure, active malignancy and the propensity for arterial injury cannot be excluded. Our human study was a retrospective analysis of clinically-indicated imaging and, therefore, the PET/CT scans were performed for clinical indications to assess for cancer rather than specifically for vascular assessments. Smaller vessels such as carotids and iliac arteries, which may be more likely to show a signal for FDG uptake, were not assessed. Patients in this cohort were most commonly treated with the ‘R-CHOP’ chemotherapy regime (rituximab, cyclophosphamide, doxorubicin, vincristine and prednisolone), which contains high-dose pulses of immunosuppression which may attenuate any inflammatory signal. Given the persisting risk of anthracycline cardiotoxicity using these chemotherapy regimens, we feel this cohort still remains a valid group for analysis^35,36^.

### Conclusions

In a large animal model, anthracycline exposure was associated with aortic fibrosis and increased intracellular vacuolization. In patients with lymphoma, anthracycline exposure was not associated with aortic inflammation assessed by ^18^F-FDG-PET/CT. Further research is required to elucidate the mechanisms of anthracycline-related vascular harm and its clinical consequences.

### Novelty & Significance

#### What is known?

- Anthracyclines are effective anti-cancer drugs but they cause arterial injury and increase arterial stiffness.
- The pathophysiologic mechanisms are poorly defined and limited to small animal and cell culture studies.
- Anthracycline-associated cardiovascular toxicities appear to share some common mechanisms with other conditions, such as hypertension and ageing, where inflammation has been established to play a role in the disease process.

#### What new information does this article contribute?

- In non-human primates, anthracycline exposure is associated with the development of arterial fibrosis and intracellular vacuolization.
- In patients with lymphoma, treatment with anthracycline exposure is not associated with large vessel inflammation when assessed by ^18^F-FDG PET/CT.
- This study suggests that inflammation may not be causally linked to anthracycline-induced aortic fibrosis and further research is required to elucidate the time course and pathophysiology of anthracycline-associated arterial fibrosis to inform mechanistically-targeted preventative strategies.

### Novelty & Significance summary

This study combined non-human primate necropsy arterial specimens with human imaging data. Anthracycline-associated arterial dysfunction has been observed in humans and has been linked to issues including heart failure but mechanistic studies have been limited to small animal models and cell lines. We sought to understand the mechanism underlying anthracycline-associated arterial injury to inform strategies to prevent this adverse effect. In our non-human primate study, animals were exposed to an anthracycline dosing regime that closely recapitulates that used in clinical practice. Necropsy specimens revealed arterial fibrosis and intracellular vacuolization in animals treated with anthracycline in comparison to controls. Anthracycline-associated arterial toxicity shares common pathways with disease processes in which inflammation plays a role in arterial remodeling. ^18^F-FDG PET/CT is the gold standard imaging modality for the assessment of large artery inflammation. Imaging analysis of ^18^F-FDG PET/CT in patients with lymphoma showed no change in aortic activity following treatment with anthracyclines suggesting that inflammation may not be responsible for aortic stiffening in patients who have received anthracyclines. Further research is required to elucidate the mechanisms of anthracycline-related vascular harm.

## Acknowledgements

None

